# Heterologous flavivirus exposure provides varying degrees of cross-protection from Zika virus in a mouse model of infection

**DOI:** 10.1101/2020.12.23.424273

**Authors:** Mariah Hassert, Stephen Scroggins, Abigail K. Coleman, Enbal Shacham, James D. Brien, Amelia K. Pinto

## Abstract

The 2015/16 Zika virus epidemic in South and Central America left the scientific community urgently trying to understand the disease and the factors which modulate Zika virus pathogenesis. Multiple other flaviviruses are endemic in areas where Zika virus emerged in 2015/16. Therefore, it is hypothesized that a key to understanding how Zika virus infection and disease progresses, is to study Zika virus infection in the context of prior flavivirus exposure. Humans and animal studies have highlighted the idea that having been previously exposed to a heterologous flavivirus may modulate the immune response to Zika virus. However, it is still unclear 1) how this impacts viral burden and pathology, and 2) the factors which correlate with the multiple metrics of disease. In this murine study, we longitudinally examine multiple factors involved in Zika disease, linking viral burden over time with increased neurological disease severity and weight loss. We show that prior heterologous flavivirus exposure with dengue virus type 2 or 3, or the vaccine strain of yellow fever, provides protection from mortality in a lethal Zika challenge. Reduction in viral burden and Zika disease in the context of prior flavivirus exposure varies depending on the infecting primary virus; with primary Zika infection being most protective from Zika challenge, followed by dengue 2, yellow fever, and dengue 3. This study demonstrates a protective effect of prior heterologous flavivirus exposure on Zika virus pathogenesis, and defines the relationship between prior flavivirus exposure and the potential for Zika virus disease.

**IMPORTANCE:** The emergence and re-emergence of various vector-borne diseases in recent years highlights the need to understand the mechanisms of protection for each pathogen. In this study, we investigated the impact of prior exposure to Zika, dengue serotypes 2, 3, and the vaccine strain of yellow fever on pathogenesis and disease outcomes in a mouse model of Zika virus infection. We found that prior exposure to a heterologous flavivirus was protective from mortality, neurological disease, weight loss, and severe viral burden during a lethal Zika challenge. Using a longitudinal study design, we were able to link multiple disease parameters including viral burden over time with neurological disease severity and weight loss in the context of heterologous infection. <u>This study demonstrates a role for heterologous flavivirus exposure in modulating flavivirus pathophysiology. Given the cyclic nature of most flavivirus outbreaks, this work will contribute to the forecasting of disease severity for future outbreaks.</u>

## INTRODUCTION

Zika virus (ZIKV) made a devastating impact when it was introduced in the Americas in 2015 and was declared a public health emergency by the World Health Organization (WHO) (1). During this ZIKV epidemic, it was reported that nearly 800,000 people in the Americas had either suspected or confirmed cases of ZIKV infection (2). While the population in the Americas was naïve to ZIKV, multiple flaviviruses including yellow fever (YFV) and the four serotypes of dengue (DENV 1-4) are endemic to the area (3, 4). With the introduction of ZIKV into the flavivirus endemic areas of the Americas the question of how and if prior flavivirus exposure could impact the course of disease with a subsequent flavivirus has become one of the most outstanding questions in flavivirus biology.

The *Flavivirus* genus consists of a number of related arthropod-borne viruses (arboviruses) which represent a substantial burden to global health and economic stability. Flaviviruses are small enveloped positive stranded RNA viruses in the family *Flaviviridae*. Following entry into susceptible cells viral replication occurs in the cytosol (5). The flavivirus genome is contained within a single open reading frame which encodes a single polyprotein. The polyprotein is cleaved into ten proteins: three structural proteins, the capsid (C), pre-membrane/membrane (prM/M), and envelope (E), as well as seven nonstructural proteins (6). As the structure and replication of flaviviruses are thought to be highly similar, our understanding of intra and extra cellular pathways of flavivirus replication cycles comes from studies of multiple different flaviviruses (Reviewed in 5, 6, 7, 8). Similarly, structural studies have long used existing knowledge of flavivirus structure to build and support the structural studies of emerging flaviviruses including ZIKV (9-11). The high degree of relatedness between flaviviruses has provided a foundation for understanding emerging flaviviruses, but has also confound epidemiological studies and diagnostics, as the high degree of relatedness makes flaviviruses more difficult to serologically distinguish flaviviruses *in vivo*.

The concern of the impact of a prior flavivirus exposure on a subsequent ZIKV infection arises from a significant precedent in the DENV literature. There is a well-established link between prior DENV exposure and enhanced disease during infection with a heterologous DENV serotype (12). With the introduction of ZIKV into the DENV and YFV endemic areas of South and Central America there is a question regarding if prior exposure to a heterologous flavivirus could enhance disease severity. Opposite to this concern is the question of whether the high degree of genetic, and structural similarities between flaviviruses (13) could afford some protection against novel circulating flaviviruses. Again, the dengue literature provides some evidence for cross-protection; with a single DENV serotype showing short-term protection against infection with heterologous serotypes (14). However, what is missing from these studies, and field studies of ZIKV infection in the Americas, is the ability to carefully monitor the infection and disease course. So while epidemiological studies have provided excellent insight into potential correlations between prior heterologous flavivirus exposure and ZIKV pathogenesis, murine models of heterologous infection are important for making more definitive links.

What we have learned from animal studies and clinical observations of flavivirus infection is that the high degree of similarity in flavivirus replication cycles, genetics and structures does not necessarily translate into similarities in cell tropism, pathogenesis, disease course, and outcomes following infection. An example of this would be the comparison between YFV and ZIKV. ZIKV infection is primarily asymptomatic in adults and children, and has shown significant tropism for the central nervous system (CNS); where YFV is primarily thought of as a hemorrhagic fever virus causing mild to severe disease in 45 percent of those infected and can have a mortality rate as high as 8 percent (WHO). So while we have been able to use the strong structural studies of flaviviruses including DENV and YFV to make rapid advancements in our understanding of ZIKV biology we have had to rely more heavily on animal models to understand the implications for disease. This is especially true in addressing the question of how a prior flavivirus infection impacts the disease course of a subsequent flavivirus infection.

Small animal models of infection have been critical for defining the correlates of protection and modeling disease for several flaviviruses, including DENV (15), YFV (16, 17), and ZIKV (18-20). In the case of ZIKV, the type 1 interferon receptor knockout mouse model (Ifnar1-/-) has been used extensively for these purposes. A few studies in murine systems have demonstrated that prior infection with a mouse adapted strain of DENV2 (strain D2S20) protects from a lethal ZIKV challenge (21), and to some degree is protective from fetal loss in maternal infection models (22). However, the impact of prior heterologous flavivirus exposure on pathogenesis still remain unclear. Namely, the consequences for viral neuroinvasion and physical indicators of neurological disease.

To address the gaps in knowledge of heterologous flavivirus exposure, we have completed a comprehensive longitudinal cross-protection study, collecting multi-parametric data to evaluate the impact of prior exposure to ZIKV, DENV2, DENV3, and the vaccine strain of YFV (YF-17D) on ZIKV neurological disease. With this study, we demonstrated that a sublethal heterologous flavivirus exposure confers varying degrees of protection from ZIKV mortality, weight loss and neurological disease. Prior exposure to either ZIKV or DENV2 were the most protective from ZIKV challenge. Exposure to YF-17D or DENV3 lessened mortality, disease severity, as well as duration of disease though some animals still succumbed to infection. Importantly, prior exposure to either ZIKV, DENV2, DENV3, or YF-17D significantly reduced viral burden in the spleen, liver, kidney, brain, and spinal cord of mice infected with ZIKV in comparison to a primary ZIKV infection. These data demonstrate a cross-protective effect of prior flavivirus exposure on ZIKV replication and disease burden. Our findings demonstrate that a subsequent flavivirus infection provide cross-protection from ZIKV disease.

## RESULTS

### DENV, ZIKV, and YFV share a significant degree of geographic and genetic overlap

Given the recent ZIKV outbreak and the high likelihood that ZIKV or another flavivirus will re-emerge and cause disease, the question of how prior heterologous flavivirus exposure influences ZIKV viral burden and disease during infection is crucial. To demonstrate the relative infection rates for ZIKV, DENV (non-serotype discriminate), YFV, or vaccination coverage with YF-17D during and following the ZIKV epidemic in the Americas, case incidence data (for ZIKV, DENV, and YFV) and YF-17D vaccine coverage data was collected from the WHO/PAHO database for South and Central American countries from 2015-2019 and displayed graphically as average annual incidence per 100,000 people over the time period of interest for each country (**Fig. 1A**). Many areas in South and Central America are considered endemic for circulation of flaviviruses including YFV, and the four serotypes of DENV (WHO/PAHO). However most people living in those regions had likely never been exposed to ZIKV prior to the 2015/2016 ZIKV outbreak (23). In 2017, as ZIKV infections continued to be documented, the northern region of Brazil was experiencing a surge of YFV cases, prompting a large vaccination campaign. Additionally, the re-emergence and dramatic increase in DENV cases in South America over the last 50 years has resulted in many regions in South America being termed “dengue hyper-endemic” (3). These most recent outbreaks are reflected in the WHO/PAHO data and demonstrate that the vast majority of South and Central American countries have reported the circulation of more than one of these flaviviruses over the past 5 years (**Fig. 1B**). The cyclic nature of the endemic flavivirus infections within this region demonstrate that even in years where disease incidence is relatively low, the prior flavivirus exposure rate within the population is relatively high (24, 25).

**Fig 1:**
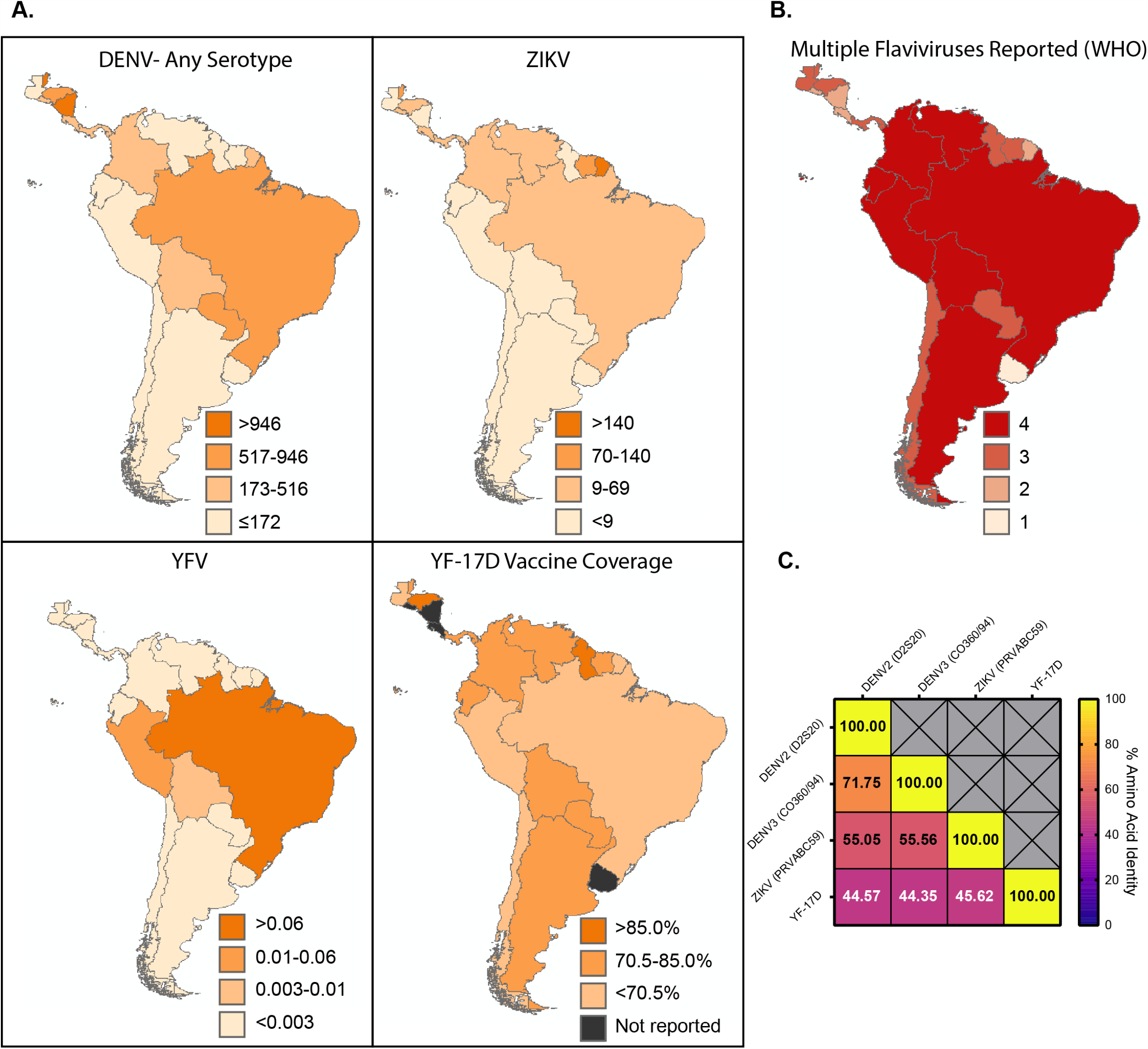
The flaviviruses contained within this study share a substantial degree of geographic and genetic overlap. **(A)** Average annual incidence rates per 100,000 people in South and Central American countries from 2015-2019. DENV, ZIKV, and YFV infections and YF-17D vaccine coverage were reported by the WHO/PAHO. Data is displayed as annual average incidence per 100,000 people in a given country for infections, or percent reported vaccine coverage for YF-17D. **(B)** The number of each of the flavivirus of interest reported in each country from 2015-2019 ranging from 1-4. **(C)** Amino acid identity of the full length polyprotein of each virus used in the current study.

There are a significant number of genetic and antigenic similarities between ZIKV, the four serotypes of DENV, and YFV (13, 26). Comparison of the amino acid identity of the full length polyproteins between ZIKV (strain PRVABC59) and DENV 2 (strain D2S20), DENV3 (strain CO360/94), and YFV (strain YF-17D) demonstrates between 44 to 71 percent identity between the viruses in various combinations (**Fig. 1C**). Based on epidemiological studies demonstrating the potential for cross-protection (25, 27) as well as extensive genetic overlap (**Fig. 1C**) (28), we hypothesized in this study that exposure to DENV, or YFV would confer some protection from ZIKV pathogenesis in a murine model.

### Prior flavivirus exposure impacts ZIKV disease progression and mortality

We first wanted to examine the cumulative effects of prior flavivirus exposure on protection from ZIKV through longitudinal sequential challenge experiments in a mouse model of infection and pathogenesis (20, 29). 4-5 week old Ifnar1-/- mice were vaccinated intravenously (IV) with 10^5^ focus forming units (FFU) of either DENV2 (mouse adapted-strain D2S20) or DENV3 (strain CO360/94). 8 week old Ifnar1-/- mice were vaccinated subcutaneously (SC) with 10^5^ FFU of the vaccine strain of YFV (YF-17D). As positive and negative controls for this experiment, 8 week old mice were vaccinated with 10^5^ FFU of ZIKV (SC) or PBS, respectively. All primary viral infections were given to mice at specific ages and at specific doses and routes of infection which were sufficient to result in infection, but not mortality (15, 17-19). Approximately 30 days following primary infection, mice were challenged with 10^5^ FFU of ZIKV IV route, which we have established in a naïve animal is a lethal route of infection, with results in 80-90% of mice succumbing to infection by day 14 (18, 19). It has been shown previously that sublethal vaccination with ZIKV confers protection from future lethal ZIKV challenges, even in intracranial challenge models (30). During ZIKV challenge, the mice were then monitored daily for 14 days for mortality and indicators of disease such as weight loss and limb weakness or paralysis. At multiple time points following ZIKV infection (day 4, 7, 14, 30), blood was collected in a subset of mice via cheek bleed and analyzed for viral burden via qRT-PCR **(Fig. 2A)**.

**Fig 2:**
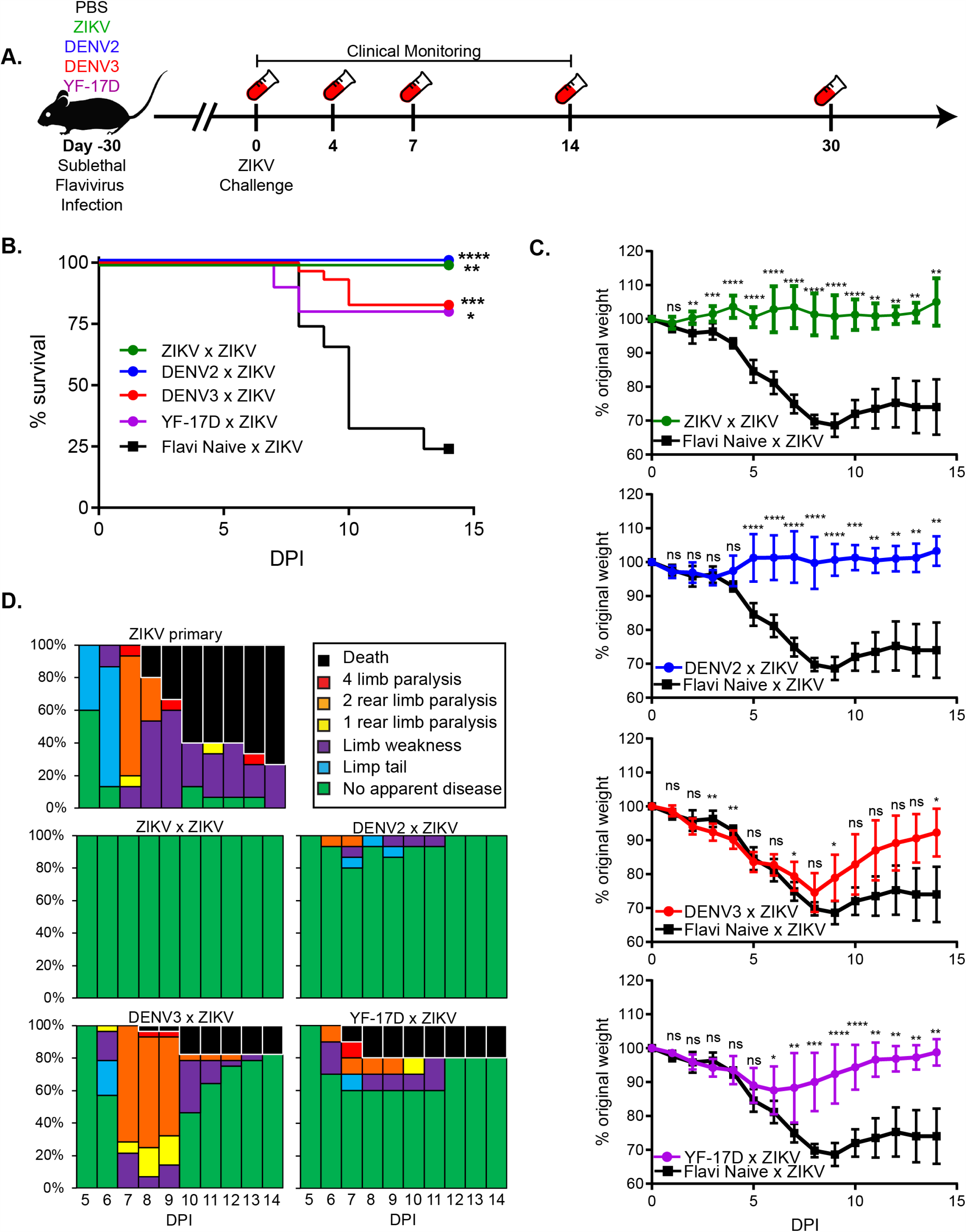
Prior flavivirus exposure leads to reduced disease severity and mortality during ZIKV challenge. (**A**) Experimental design. Ifnar1-/- mice were sublethally infected with either ZIKV (n=8), DENV2 (n=15), DENV3 (n=30), or YF-17D (n=10) or PBS as a flavivirus naïve control (n=11). 30 days following primary infection, mice were challenged with ZIKV by IV administration. For 14 days following ZIKV challenge, mice were monitored for indicators of neurological disease as previously described (18, 19), weight loss, and mortality. At day 4, 7, 14, and 30 post ZIKV challenge, blood was collected to measure viremia by qRT-PCR. Data is a compilation of 4 independent experiments with at least 8 animals per group. (**B**) Survival of Ifnar1-/- mice with or without prior flavivirus exposure during IV ZIKV challenge. Survival rates of mice with prior ZIKV exposure (ZIKV × ZIKV), prior DENV2 exposure (DENV2 × ZIKV), prior DENV3 exposure (DENV3 × ZIKV), or prior YF-17D exposure (YF-17D × ZIKV) were significantly greater than mice with no prior flavivirus exposure during ZIKV infection as determined by Mantel-Cox test (*p=0.0332, **p=0.0021, ***p=0.0002, ***p<0.0001). (**C**) Weight loss during ZIKV challenge. As a measure of disease burden, mice were weighed daily for 14 days post ZIKV challenge. Weight change is displayed by normalizing relative to the starting weight of each animal on the day of ZIKV challenge. Compared to mice that were flavivirus naïve at the time of ZIKV challenge, mice with prior flavivirus exposure displayed less weight loss over time as determined by Two-Way ANOVA with Dunnett’s post hoc analysis (*p=0.0332, **p=0.0021, ***p=0.0002, ***p<0.0001). (**D**) Neurological indicators of ZIKV disease. Mice were evaluated daily for sequela associated with ZIKV infection and graphed as a percentage of the total number of mice per group. Disease onset began on day 5 for mice with no prior flavivirus exposure and lasted or progressed for at least 9 days. Mice with prior ZIKV exposure displayed no indicators of neurological dysfunction during ZIKV challenge. Mice with prior DENV2 exposure experienced a low frequency of neurological disease between days 5-11. Mice with prior DENV3 exposure experienced a high frequency of neurological disease though most of the mice recovered by day 14. Mice with prior YF-17D exposure displayed a moderate frequency of neurological disease which most recovered from by day 12.

Consistent with previous literature (30), mice with prior ZIKV exposure (ZIKV × ZIKV) were completely protected from overt ZIKV induced morbidity and mortality (**Fig. 2B-D**,) compared to mice with no prior flavivirus exposure (Flavi naïve × ZIKV), which lost a significant amount of weight (**Fig. 2C**), and all suffered from ZIKV-induced neurological indicators of disease including flaccid tail, limb weakness, hind limb paralysis, or complete limb weakness or paralysis (**Fig. 2D**). Approximately 75% of mice challenged with ZIKV in the absence of prior flavivirus exposure eventually succumbed to infection, consistent with our previous reports (18, 19) (**Fig. 2B**).

Upon infection with ZIKV, mice with prior heterologous flavivirus exposure had significantly reduced mortality relative to flavivirus naïve mice infected with ZIKV. 100% of mice with prior DENV2 exposure (DENV2 × ZIKV), 80% of mice with prior DENV3 exposure (DENV3 × ZIKV), and 80% of mice withYF-17D (YF-17D × ZIKV) exposure survived ZIKV challenge (**Fig. 2B**). Moreover, having prior heterologous flavivirus exposure conferred varying degrees of protection from ZIKV-induced morbidity in this model. In the case of DENV2 and YF-17D exposure, these mice lost significantly less weight than the flavivirus naïve mice during ZIKV challenge (Flavi naïve × ZIKV) (**Fig. 2C**). Substantially fewer mice in these groups displayed any signs of neurological impairment, and the duration of disease appeared to be shorter compared to the Flavi naïve × ZIKV group (**Fig. 2D**). In the case of prior DENV3 exposure (DENV3 × ZIKV), partial protection from ZIKV was conferred, as indicated by only a mild reduction in weight loss relative to the flavi naïve × ZIKV group, during ZIKV challenge (**Fig. 2C**). These mice were clearly not as resistant to ZIKV induced weight loss as the other groups with prior flavivirus exposure, and all mice within this group displayed some signs neurological involvement (**Fig. 2D**). Overall, this data shows that prior heterologous flavivirus exposure is generally protective from ZIKV induced morbidity and mortality in the Ifnar1-/- mouse model, though varying degrees of cross-protection can be observed depending upon the primary infecting virus.

### ZIKV viremia over time is differentially influenced by prior flavivirus exposure

Throughout the course of this longitudinal study (**Fig. 2A**), blood was collected at multiple timepoints post ZIKV exposure and analyzed by qRT-PCR to assess the impact of heterologous flavivirus exposure on ZIKV viremia. Consistent with our previous studies using this model (18, 19), during a primary ZIKV infection (Flavi naïve × ZIKV), viral RNA is detectable in the blood by day 4 post-infection, and trends in a downward trajectory over time, however, the virus is not cleared even by day 30 in these animals (**Fig. 3**). Sublethal vaccination with ZIKV prior to ZIKV challenge results in a substantial reduction in in viremia (relative to mice with no prior flavivirus exposure) starting as early as 4 days post challenge, and continuing throughout the course of the experiment (**Fig. 3A**). This is consistent with previous reports demonstrating the protective capacity of ZIKV immunity upon rechallenge with ZIKV (30-32).

**Fig 3:**
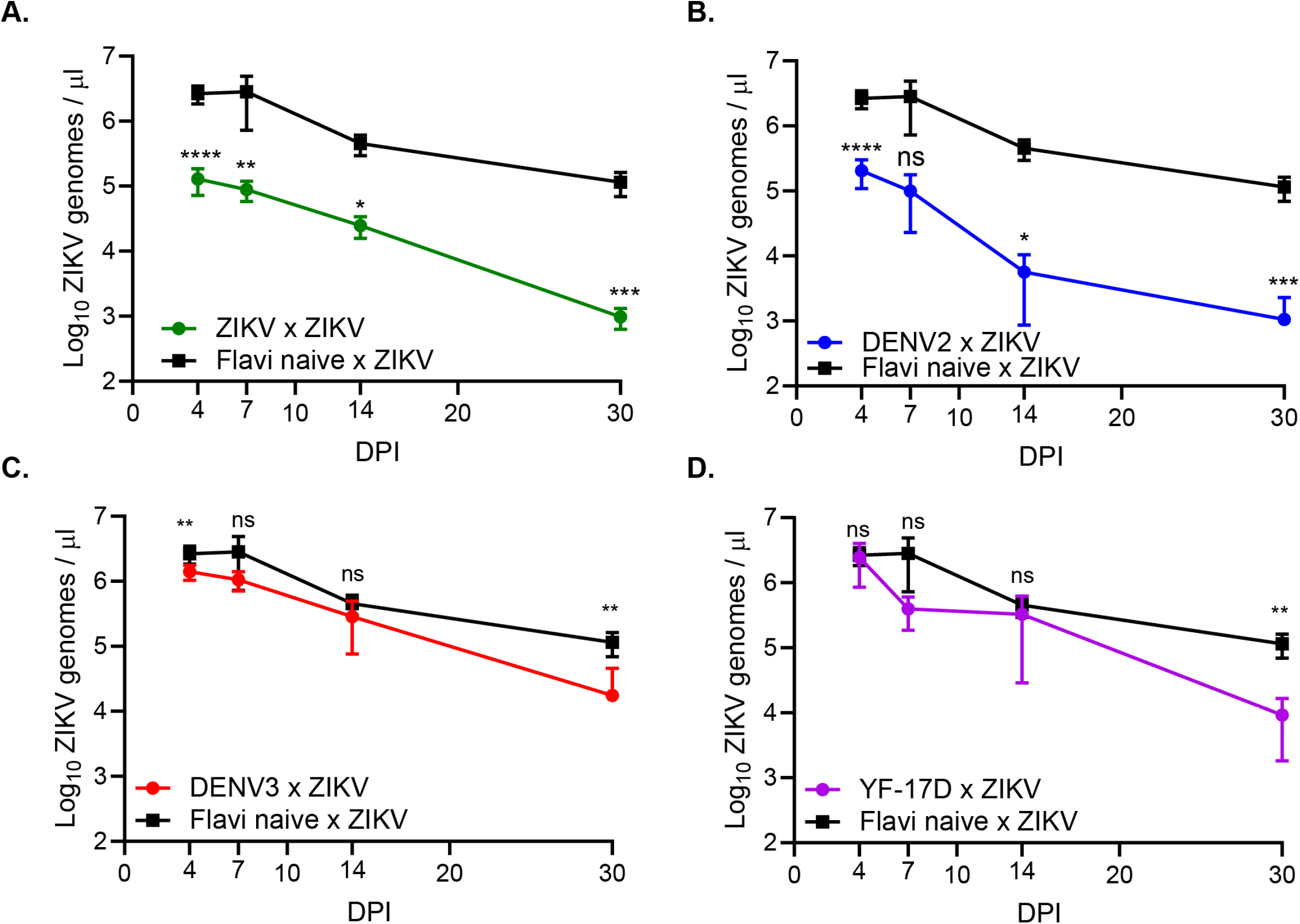
ZIKV viremia over time following heterologous challenge. Following ZIKV challenge, blood was collected via submandibular bleed on days 4, 7, 14, and 30 to evaluate viremia over time by qRT-PCR. (**A**) ZIKV viremia of homologously primed and boosted (ZIKV × ZIKV) mice compared to mice with no prior flavivirus exposure during ZIKV challenge. (**B**) ZIKV viremia of DENV2 immune mice during ZIKV challenge (DENV2 × ZIKV) mice compared to mice with no prior flavivirus exposure during ZIKV challenge. (**C**) ZIKV viremia of DENV3 immune mice during ZIKV challenge (DENV3 × ZIKV) mice compared to mice with no prior flavivirus exposure during ZIKV challenge. (**D**) ZIKV viremia of YF-17D vaccinated mice during ZIKV challenge (YF-17D × ZIKV) mice compared to mice with no prior flavivirus exposure during ZIKV challenge. Statistically significant differences in viremia over time over time was determined by Two-Way ANOVA with Dunnett’s post hoc analysis (*p=0.0332, **p=0.0021, ***p=0.0002, ***p<0.0001).

Similar to the influence of prior heterologous flavivirus infection on ZIKV neurological disease and weight loss, viremia was variable dependent upon the virus that was given upon primary infection. In mice with prior DENV2 exposure, ZIKV viremia was statistically lower throughout the course of infection, with the exception of day 7, relative to mice with no prior flavivirus exposure (**Fig. 3B**). However, in mice with prior DENV3 or YF-17D exposure, over time there was a statistically significant reduction in ZIKV viremia during heterologous secondary infection relative to a primary infection only at days 4 and 30 for mice with prior DENV3 exposure and day 30 for mice with prior YF-17D exposure **(Fig. 3C-D)**.

### Global ZIKV viral burden is reduced with prior flavivirus exposure

While we observed a reduction in ZIKV viremia over time in mice that had been previously exposed to another flavivirus, it was unclear how heterologous flavivirus exposure could influence viral burden in tissues both within the peripheral organs and central nervous system (CNS). ZIKV invasion of the CNS appears to be a crucial factor in disease pathology within murine models of infection, as well as in human disease (19, 20, 30, 33), though the mechanism that drives this is not fully understood. To determine the impact of prior heterologous flavivirus exposure on the viral burden in various organs during ZIKV infection, we sublethally infected mice with either ZIKV, DENV2, DENV3, YF-17D, or a vehicle control. 30 days post infection, the mice were challenged IV with ZIKV as described in Figure 2A. At day 4 and 8 post ZIKV challenge, the spleen, liver, kidneys, brains, and spinal cords were harvested and homogenized. RNA was extracted from these homogenates and ZIKV viral burden was determined by qRT-PCR (**Fig. 4**).

**Fig 4:**
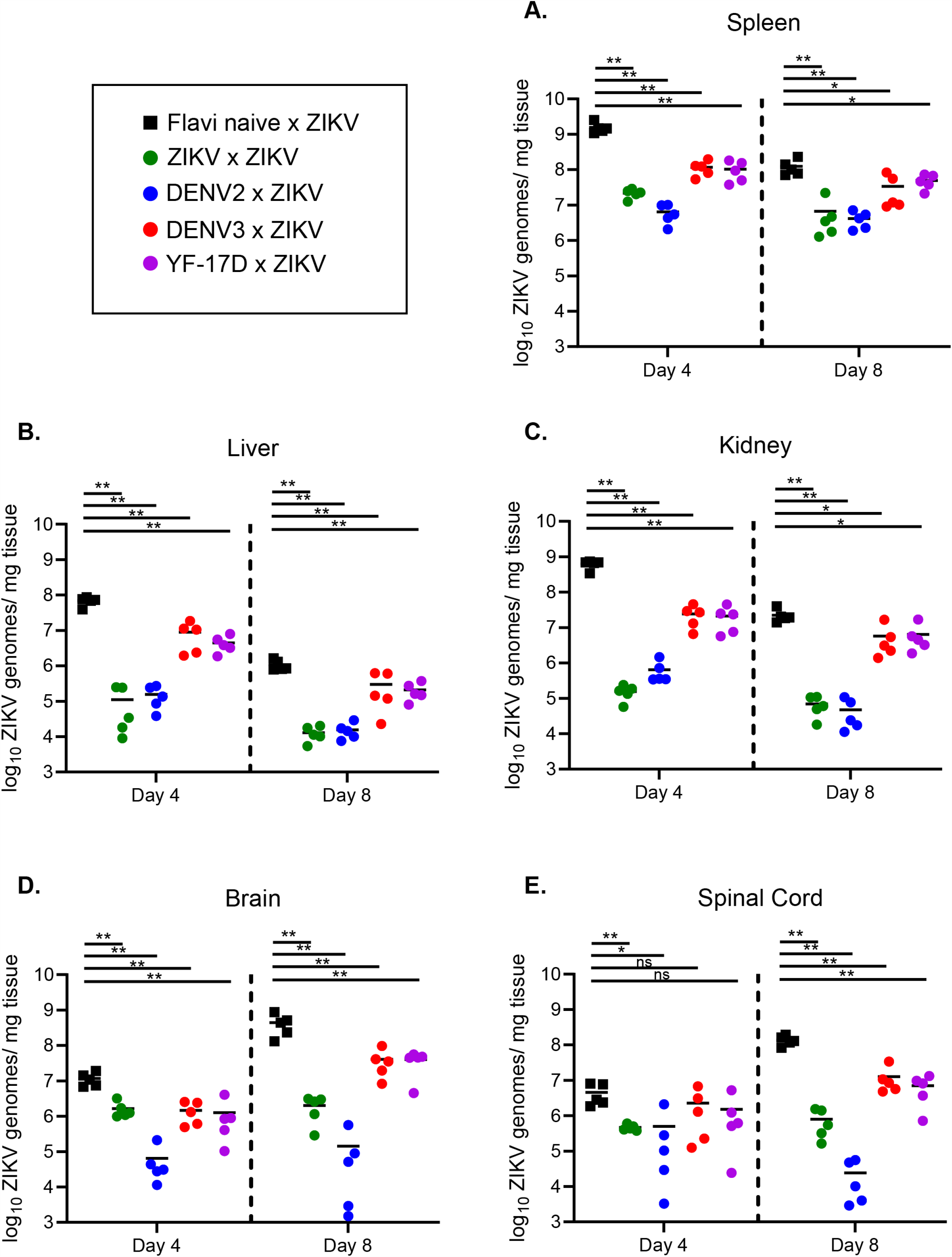
ZIKV viral burden in the peripheral organs and CNS is reduced with prior flavivirus exposure. Ifnar1-/- mice were sublethally infected with either ZIKV (n=10), DENV2 (n=10), DENV3 (n=10), or YF-17D (n=10) or PBS as a flavivirus naïve control (n=10). 30 days following primary infection, mice were challenged with ZIKV by IV administration. At days 4 and 8 post ZIKV challenge (n=5 mice per group, per day), mice were euthanized, perfused with PBS, and organs were weighed and snap frozen. RNA was extracted and qRT-PCR was performed to measure viral burden in the (**A**) Spleen, (**B**) Liver, (**C**) Kidney, (**D**) Brain, and (**E**) Spinal Cord. Data is displayed as Log_10_ ZIKV genome copies per mg of tissue. Statistical significance was determined by Mann-Whitney test (*p=0.0332, **p=0.0021, ***p=0.0002, ***p<0.0001).

Similar to previous reports of primary ZIKV infection in the Ifnar1-/- model, (19, 20, 29) day 4 appears to be the peak in viral burden in most peripheral organs (**Fig. 4A-C**), while day 8 is the peak for viral burden in CNS tissues (**Fig. 4D-E**). By the time of peak neurological pathology in mice with no prior flavivirus exposure (day 8-9), the virus has invaded both the brain and spinal cord and replicated to high titers. As expected, mice with prior ZIKV exposure display significantly reduced viral burden in the spleen (**Fig. 4A**), liver (**Fig. 4B**), kidney (**Fig. 4C**), brain (**Fig. 4D**), and spinal cord (**Fig. 4E**) relative to flavi naïve × ZIKV mice on both days 4 and 8. However, we find it important to note that this significant reduction does not appear to be completely sterilizing as viral genomes are still being detected on both days in all tissues. We have previously reported that ZIKV is a persistent infection in the Ifnar1-/- model (19), which could potentially be the reason for this observation.

In the case of mice with prior heterologous flavivirus exposure (DENV2, DENV3, or YF-17D), we saw a significant reduction in ZIKV viral burden in peripheral tissues on both days 4 and 8 relative to the flavi naïve × ZIKV group (**Fig. 4A-C**). The most drastic of these reductions came from mice with prior DENV2 exposure, which seems to trend well with the observation of this being the group of heterologously challenged mice with the least severe disease (**Fig. 2**). On day 4 post infection, we observed a significant reduction in ZIKV viral load in the brains of all groups of mice with prior flavivirus exposure (**Fig. 4D**). However, in the spinal cord, only heterologously challenged mice with prior DENV2 exposure had statistically significantly reduced viral loads on day 4 (**Fig. 4E**). By day 8, the peak in disease burden and viral burden in the CNS, all groups of heterologously challenged mice displayed reduced viral load in both the brain and spinal cord (**Fig. 4D-E**). Overall, this data demonstrates that prior heterologous flavivirus exposure impacts the outcome of ZIKV challenge by a global reduction in viral burden on days 4 and 8 post infection.

### Defining the relationships of multiple disease metrics during heterologous ZIKV challenge

Disease metrics assessed during ZIKV infection in mouse models are diverse. Studies using these models (including our own), have quantified disease using non-invasive techniques such as weight loss, neurological disease assessment, mortality, viremia, and viral shedding in the urine as well as more invasive techniques such as viral burden in multiple target organs, fetal resorption and loss, and neuroinvasion, inflammation, and apoptosis histologically (18-20, 22, 29, 33-37). However, it is unclear how each disease parameter in this complex system are related to one another and whether this is influenced by prior heterologous flavivirus exposure. Therefore, we generated a data bank using longitudinal data points from variables measured in **Figures. 2 and 3**, from each individual mouse. We used this data to determine correlative relationships between various metrics of disease including peak percent weight loss, day of peak weight loss, peak disease score, day of peak disease, number of days of disease, and viremia on days 4, 7, 14, and 30 post ZIKV infection by linear regression and Pearson correlation (**Fig. 5A**). The peak weight loss percentage was determined by normalizing the starting weights of each animal at day 0 to 100% and assessing weight daily and noting the peak percentage of weight lost through the course of infection. The peak disease score was determined by tracking neurological indicators as previously described, daily following ZIKV infection (18, 19). Each indicator of disease was assigned a number from 0 to 6, indicative of severity (0=no disease, 1=limp tail, 2=hind limb weakness, 3=single hind limb paralysis, 4=bilateral hind limb paralysis, 5=full body weakness/paralysis, and 6=death). The peak day of disease corresponded to the day of peak neurological disease. The number of disease days was determined by counting each day for each mouse that the disease score was above 0.

**Fig 5:**
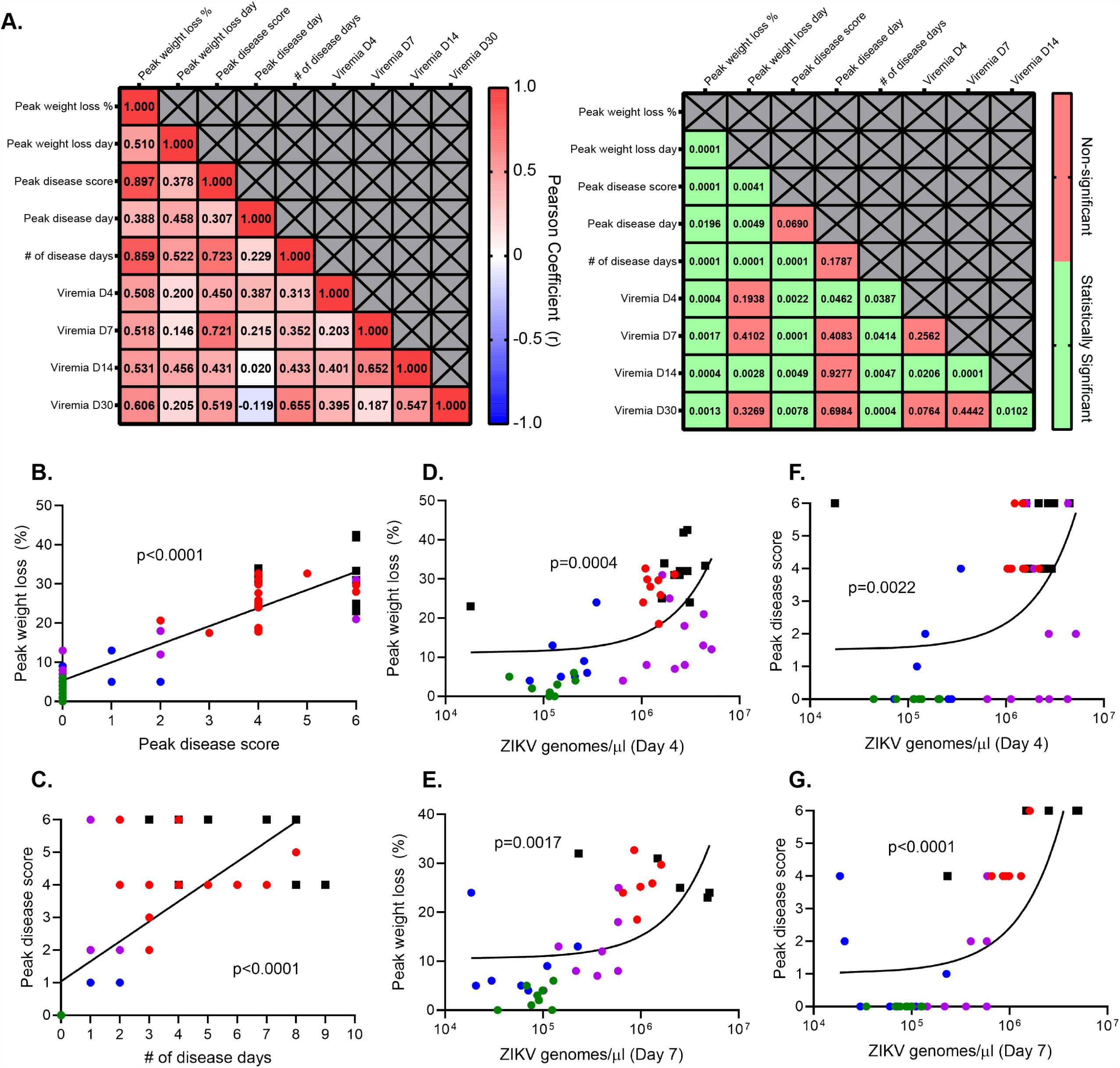
Multiple metrics of ZIKV infection and disease burden are correlated. The data collected from figures 1 and 2 was used to generate a data bank and analyzed using linear regression and Pearson correlation analysis to define the relationships of multiple disease metrics during heterologous ZIKV challenge. (**A**) A correlation matrix was generated, which displays the Pearson coefficient (r) demonstrating the strength and directionality of the correlation between each variable (left) and the p value demonstrating statistical significance of each correlation (right). (**B**) Linear regression and correlation between the peak weight loss of each animal and peak in disease score. Mice were weighed daily and observed for clinical signs of disease. Each phenotype was assigned a number from 0-6 (0=no disease, 1=limp tail, 2=hind limb weakness, 3=single hind limb paralysis, 4=bilateral hind limb paralysis, 5= full body weakness/paralysis, and 6=death). (**C**) Linear regression and correlation between peak disease score (as described in Figure 4B) and the number of days a given animal experienced clinical signs of neurological disease. (**D**) Linear regression and correlation between ZIKV genomes detected in the blood by qRT-PCR on day 4 post ZIKV infection and peak weight loss. (**E**) Linear regression and correlation between ZIKV genomes detected in the blood by qRT-PCR on day 7 post ZIKV infection and peak weight loss. (**F**) Linear regression and correlation between ZIKV genomes detected in the blood by qRT-PCR on day 4 post ZIKV infection and peak disease score. (**G**) Linear regression and correlation between ZIKV genomes detected in the blood by qRT-PCR on day 7 post ZIKV infection and peak disease score.

Of the 36 bivariate permutations, we identified 25 statistically significant correlative interactions. The strength and directionality of each correlation is indicated by the Pearson coefficient (r) and the statistical significance of each correlation is indicated by the p value (**Fig. 5A**). From these data, we identified several interactions of particular interest (**Fig. 5B-G**). The strongest correlation resulted when comparing the peak neurological disease score from each mouse to the peak percent weight lost (r=0.8973, p<0.0001) (**Fig. 5B**). As one might expect, as the peak neurological disease score identified in each animal increased, so did the amount of weight lost. Importantly, each infection group (flavi naïve × ZIKV, DENV2 × ZIKV, DENV3 × ZIKV, YF-17D × ZIKV, and ZIKV × ZIKV) generally clustered together and along the pattern determined by linear regression. This would suggest similarities in the biological characteristics of these groups. We were also interested in the relationship between the number of days each infected animal displayed a neurological disease phenotype, and the severity of the disease phenotype (**Fig. 5C**). We found that the peak disease score and the number of measured disease days strongly positively correlated with one another (r=0.7234, p<0.0001); that is in general, the more severe the disease phenotype, the longer it would take to resolve.

We were particularly interested in the relationship between viremia and less invasive metrics of disease burden such as weight loss and disease score. When comparing viral burden on day 4 or day 7 to peak percentage weight loss, (**Fig. 5D-E**), respectively) we found a statistically significant positive correlation (r=0.5078 and 0.5184, p=0.0004 and 0.0017, respectively). This was also true when comparing viral burden on day 4 or day 7 to peak disease score (**Fig. 5F-G**) (r=0.45 and 0.7213 and p=0.022 and <0.0001, respectively). That is, with increased viremia on these days came increased weight loss and neurological disease. The correlative analyses associating disease severity and viral load at day 4 are of particular interest due to the timing. As demonstrated in **Fig. 2**, neurological indicators of disease are not overtly detectable until day 5 post infection and typically do not peak until days 7-9 (**Fig. 2D**) and the most significant drop in weight also occurs from days 7-9 (**Fig. 2C**). However, using the disease metric of early viremia on day 4 it is clear that information could be used in linear regression analysis to predict the severity and outcome of infection days earlier than the occurrence of overt disease (**Fig. 5D and 5F**). Overall, these data demonstrate the bivariate interactions between various minimally invasive metrics that are commonly used to assess ZIKV disease burden in mouse models of infection and importantly, that in the context of heterologous infection that these correlations are still appropriate and comparable.

## DISCUSSION

Increased globalization, deforestation, climate change, and the lack of effective vaccines has resulted in most of the world’s population being at risk for infection with multiple flaviviruses (38). There is no vaccine available for ZIKV and current vaccines for flaviviruses including the yellow fever vaccine, while highly effective, have not prevented outbreaks from these highly prevalent arboviruses. While the current number of ZIKV cases in the Americas has dropped significantly compared to 2016, based on the infection cycles of similar flaviviruses, is believed that ZIKV will follow a similar cyclical pattern of emergence and re-emergence (39, 40). Therefore, it is highly likely that ZIKV is in an inter-epidemic period and will re-emerge and continue to spread throughout the Americas, as has been seen with both DENV 1-4, and YFV.

The influence of prior flavivirus exposure on ZIKV protection and pathogenesis remains an important question. Epidemiological studies do provide some insight into these competing concerns. A study in Brazil comparing YFV vaccination coverage with incidence rates of ZIKV associated microcephaly found that Northeast Brazil, which had the highest incidence of ZIKV associated microcephaly also had relatively low YFV vaccination rates, suggesting that a lack of YFV vaccination left that population without a cross-protective response and therefore susceptible to ZIKV morbidity (27). Moreover, the Eva Harris group has examined the relationship between prior DENV exposure and the incidence of asymptomatic ZIKV infection in a pediatric cohort in Nicaragua, finding that children with prior DENV infection had lower rates of symptomatic ZIKV infection, again suggesting ZIKV cross-protection mediated by previous DENV exposure (25). However, drawing clear causal links between previous exposure and infection outcomes are a challenge in human populations for several reasons in this case. These co-circulating flaviviruses share antigenic similarities, which can confound many serologically based diagnostic tests, which makes confirming records of the natural history of infection particularly challenging. Additionally, the length of time between exposures of heterologous serotypes of DENV plays a major role in whether increased incidence of enhanced pathogenesis or cross-protection occurs (12, 41, 42).

In this study, we challenged the hypothesis that prior heterologous flavivirus exposure to DENV serotypes 2 or 3, or YF-17D would confer cross-protection from ZIKV in a mouse model. We ultimately showed that a sublethal heterologous flavivirus exposure confers varying degrees of protection from ZIKV mortality, weight loss and neurological disease. Prior exposure to ZIKV or DENV2 were the most protective from ZIKV challenge with no mice succumbing to infection and few, if any displaying any signs of neurological disease and weight loss. Exposure to YF-17D or DENV3 lessened mortality, disease severity, and viral burden though some animals still succumbed to infection. Importantly, prior exposure to either ZIKV, DENV2, DENV3, or YF-17D significantly reduced viral burden in the spleen, liver, kidney, brain, and spinal cord of mice infected with ZIKV. These data demonstrate a cross-protective effect of prior flavivirus exposure on ZIKV replication and disease burden.

Murine models of ZIKV infection and heterologous flavivirus challenge have used diverse metrics for quantifying disease burden (18-20, 22, 29, 33-37). Until now, the relationship of many of these variables has not been evaluated. In this longitudinal heterologous challenge experiment, we performed linear regression and correlation analysis to determine the relationship between multiple variables in individual mice including peak weight loss, day of peak weight loss, peak disease score, day of peak disease score, number of disease days, viremia on day 4, viremia on day 7, viremia on day 14, and viremia on day 30. Using this analysis, we identified 25 statistically significant correlative interactions. Importantly, we found that viral burden on day 4 strongly correlated with the peak weight loss and peak disease score that an animal would eventually experience (typically on day 7-9 post infection). This allows for the possibility of using early viremia data as a predictor of severe disease outcomes using linear regression analysis that can also be applied in the context of heterologous infection scenarios.

Ultimately, these data provide additional evidence of the cross-protective effect of prior heterologous flavivirus exposure on ZIKV disease. These findings are important given that the majority of the world is at risk of flavivirus exposure and many regions are endemic for multiple flaviviruses. Addressing this is not only important for being able to predict the outcome of ZIKV exposure in flavivirus endemic areas but will support efforts to generate a pan-flavivirus vaccine. While this study provides significant insight into cross-protection from ZIKV, additional studies are desperately needed to understand the mechanism behind this. Studies such as these will be essential to control these significant public health threats.

## MATERIALS AND METHODS

### Ethics statement

All animal studies were done in accordance with the Guide for Care and Use of Laboratory Animals of the National Institutes of Health and approved by the Saint Louis University Animal Care and Use Committee (IACUC protocol #2667).

### Viruses and cells

P_0_ stocks of ZIKV, strain PRVABC59 (GenBank accession KU501215.1) and YFV, strain 17D (Genbank accession X03700) were acquired from Biodefense and Emerging Infection (BEI) Resources. Each virus was passaged in African green monkey kidney epithelial cells (Vero-WHO) that were purchased from American Type Culture Collection (ATCC CCL-81). The supernatants of these cultures were clarified of cellular debris by centrifugation at 3,500 RPM prior to being aliquoted and frozen at −80°C. DENV2, strain D2S20, (GenBank accession HQ891024) was a kind gift from Dr. Michael Diamond (43). DENV3, strain CO360/94 (GenBank accession AY923865) was obtained from ATCC. Both DENV2 and DENV3 were grown in C6/36 *Aedes albopictus* cells (ATCC CRL-1660). At the time of harvest, the media supernatant was clarified of cellular debris by centrifugation at 3,500 RPM. Each virus was then concentrated by ultracentrifugation at 30,000 RPM over a 25% glycerol cushion, before being aliquoted and frozen at −80°C (15). The infectious titer of each viral stock was quantified by focus forming assay (FFA) as previously described (34). Briefly, a 90% confluent monolayer or Vero-WHO cells were plated in a 96-well flat bottom plate. Serial dilutions of each viral stock were added to each well for 1 hour, prior to the addition of a methyl cellulose layer, to restrict lateral spread of the virus. After 48 hours (ZIKV and YF-17D) or 72 hours (DENV2 and DENV3), the cells were fixed and permeablized. The cells were incubated with a flavivirus cross-reactive monoclonal primary antibody (4G2) for 1 hour at room temperature, washed, incubated with a horse-radish peroxidase conjugated anti-mouse secondary antibody for 1 hour at room temperature, and washed. Foci of infected cells were visualized and quantified following the addition of True-Blue peroxidase substrate.

### Mice and infections

Interferon αβ receptor 1 knockout (Ifnar1-/-) mice were purchased from Jackson Laboratories. They were bred and maintained at Saint Louis University in a specific-pathogen-free mouse facility. To achieve a primary infection, at 4-5 weeks of age, equal ratios of male and female mice were administered a sublethal intravenous (IV) challenge of either DENV2 (10^5^ FFU) or DENV3 (10^5^ FFU). To generate mice with prior ZIKV or YF-17D exposure, 8 week old mice at equal ratios of male and female animals, were administered a sublethal subcutaneous (SC) challenge of either ZIKV (10^5^ FFU) or YF-17D (10^5^ FFU). The viral doses, routes of administration, and ages of mice were deliberately chosen based on optimized dosing experiments in our lab known to induce detectable viral replication and immune responses, but not cause mortality (15, 17-19). As a flavivirus naïve group, 8-week old littermate controls were administered PBS. At least 30 days following primary challenge, mice were administered an IV ZIKV challenge (10^5^ FFU), previously demonstrated by our lab to result in severe neurological sequela and weight loss in 100% of adult Ifnar1-/- mice with no prior flavivirus exposure with 80-100% ultimately succumbing to infection (18, 19). Following ZIKV challenge, mice were monitored daily for 14 days for weight loss, indicators of neurological disease, and mortality. Approximately 150 μl of blood was collected longitudinally from each mouse at day 0, 4, 7, 14, and 30 to monitor peripheral viral burden.

### Measurement of viral burden

For longitudinal studies, whole blood was collected by cheek bleed into EDTA coated tubes. 50 μl of blood was transferred to RNAsol BD reagent and RNA was extracted according to the manufacturer’s instructions. For studies evaluating global viral burden, on day 4 and 8 post ZIKV infection, mice were administered a lethal cocktail of ketamine/xylazine before intracardiac perfusion with 20 ml of PBS. The spleen, liver, kidney, brain and spinal cord were collected from each mouse and snap frozen in a dry ice bath. Organs were weighed and homogenized in DMEM using a BeadMill 24 from Fisher Scientific. RNA was extracted from 100 μl of homogenate using TriReagent RT according to manufacturer’s instructions. ZIKV RNA was quantified by qRT-PCR using a Prime-Time primer-probe set: Forward-CCGCTGCCCAACACAAG, Reverse-CCACTAACGTTCTTTTGCAGACAT, Probe-AGCCTACCTTGACAAGCAGTCAGACACTCAA.

### Statistical analysis

For DENV, ZIKV, YFV, and YF-17D incidence maps, incidence data and vaccine coverage data was collected for South and Central American countries from the WHO/PAHO for the years 2015-2019 and displayed as the annual average number of cases per 100,000 individuals using the spatial data program Geoda (44). Amino acid identity for each flavivirus of interest was determined by performing a global alignment allowing for free ends using a Blosum62 cost matrix in the software Geneious. Statistical analyses for in vivo studies were performed using Graph Pad Prism. Statistical differences in survival were determined using a Mantel-Cox test. Differences in weight loss and viral burden over time were determined using a Two-Way ANOVA with post-hoc analysis. Viral burden in various organs at points of organ harvest were determined by Mann-Whitney test. Correlative analysis was performed using linear regression analysis and a two-tailed Pearson analysis.

### Data Availability

https://apps.who.int/gho/data/view.main.1540_50?lang=en, https://apps.who.int/immunization_monitoring/globalsummary/timeseries/tscoverageyfv.html, https://www.paho.org/data/index.php/en/mnu-topics/zika/524-zika-weekly-en.html https://www.paho.org/data/index.php/en/mnu-topics/indicadores-dengue-en/dengue-nacional-en/252-dengue-pais-ano-en.html

## Author Contributions

Conceptualization-MH, JDB, AKP. Data curation-MH, SS. Formal Analysis-MH. Investigation-MH. Methodology-MH, JDB, AKP, SS. Funding Acquisition-MH, JDB, AKP. Resources-AKC, SS, ES, TLS. Visualization-MH, SS. Writing (original draft)-MH, JDB, AKP. Writing (review and editing)-MH, SS, TLS, AKC, ES, JDB, AKP. Project administration-JDB, AKP.

